# Spatial Transcriptomic Analysis Identifies Epithelium-Macrophage Crosstalk in Endometriotic Lesions

**DOI:** 10.1101/2024.03.23.586434

**Authors:** Gregory W. Burns, Zhen Fu, Erin L. Vegter, Zachary B. Madaj, Erin Greaves, Idhaliz Flores, Asgerally T. Fazleabas

## Abstract

The mechanisms underlying the pathophysiology of endometriosis, characterized by the presence of endometrium-like tissue outside the uterus, remain poorly understood. This study aimed to identify cell type-specific gene expression changes in superficial peritoneal endometriotic lesions and elucidate the crosstalk among the stroma, epithelium, and macrophages compared to patient-matched eutopic endometrium. Surprisingly, comparison between lesions and eutopic endometrium revealed transcriptional similarities, indicating minimal alterations in the sub-epithelial stroma and epithelium of lesions. Spatial transcriptomics highlighted increased signaling between the lesion epithelium and macrophages, emphasizing the role of the epithelium in driving lesion inflammation. We propose that the superficial endometriotic lesion epithelium orchestrates inflammatory signaling and promotes a pro-repair phenotype in macrophages, providing a new role for Complement 3 in lesion pathobiology. This study underscores the significance of considering spatial context and cellular interactions in uncovering mechanisms governing disease in endometriotic lesions.

## Introduction

Endometriosis affects around 10% of reproductive age women and is defined by the presence of endometrium-like tissue growing outside of the uterus [1–3]. The disease is clinically associated with life-altering pain and infertility and up to 50% of infertile women are diagnosed with endometriosis [4, 5]. Endometriotic lesions are estrogen-dependent and the only effective non-surgical treatment is suppression of ovarian function to limit estrogen production. Endometriotic lesions in the peritoneal cavity are classified by location with lesions present on the visceral or parietal peritoneum termed peritoneal lesions and those on the ovary termed endometriomas. Peritoneal lesions are further separated into superficial or deep infiltrating according to the invasion depth of underlying tissue with deep infiltrating lesions invading more than 5 mm [6, 7]. Superficial peritoneal endometriotic lesions likely develop from endometrial tissue fragments that are refluxed into the peritoneal cavity during menstruation [8, 9]. This hypothesis is supported by the baboon model of endometriosis in which menstrual tissue inoculated into the peritoneal cavity results in endometriotic lesions and disease that persists for at least 15 months [10–12]. Endometriosis can also be induced in mice with tissue collected from an artificial menstruation cycle [13, 14].

Superficial peritoneal lesions generally consist of endometrium-like epithelium and stroma, fibroblasts that express smooth muscle actin (ACTA2), and immune cells [15]. The presence of ACTA2+ fibroblasts is positively correlated with fibrotic lesions where the presence of epithelium is less common, in contrast to lesions of endometrium-like stroma and epithelium [16, 17]. During lesion development, activation of pro-inflammatory immune populations, specifically macrophages, provide a hospitable inflammatory environment to protect the ectopic endometrial tissue [18]. Previous bulk and single-cell RNA-sequencing analyses have uncovered thousands of differentially expressed genes (DEG) in endometriotic lesions, notably including inflammation- and immune-related genes [19–23]. While bulk RNA-sequencing offers comprehensive data, it lacks the ability to discern information about individual cell types. Conversely, single-cell RNA-sequencing provides insights into cell type-specific expression but relies on post-hoc cell type identification using known markers and lacks spatial context. The emergence of spatial transcriptomics has addressed these limitations by providing a powerful tool to understand the spatial context of gene expression within tissues, particularly when coupled with canonical cell type immunostaining. Two of the most prominent platforms for spatial transcriptomics, NanoString GeoMx and 10X Visium, offer distinct advantages and limitations depending on experimental design. In this study, we chose the GeoMx platform based on the ability to provide cell type specific data from formalin fixed paraffin embedded (FFPE) tissues with high specificity based on antibody-based fluorescent segmentation [24–26]. Our aim was to determine the genes and pathways altered in the glandular epithelium, stroma, and macrophages within endometriotic lesions compared to matched endometrium with spatial transcriptomics. We hypothesized that gene expression changes in superficial peritoneal endometriotic lesions would manifest in a cell type-specific manner, with disruption of the crosstalk among the stroma, epithelium, and macrophages being evident compared to patient-matched eutopic endometrium.

## Results

### Spatial Transcriptomics of Human Superficial Peritoneal Lesions

We sought to determine the contributions of epithelium, stroma, and myeloid cell compartments of endometriotic lesions to altered genes and pathways in endometriotic lesions by spatial transcriptomic analyses. Superficial peritoneal lesions and eutopic uterine tissue were collected from women in the secretory phase of the menstrual cycle. Matched lesion and eutopic tissues from five women (n = 10) were selected based on the histological presence of endometrial gland-like structures in lesions confirmed by trichrome staining (Figure 1). The average age of the patients was 42 (Table 1, SD = 6.1). Tissues were stained for pan-cytokeratin, smooth muscle actin, and CD68 to identify epithelium (pan-cytokeratin+), stroma (ACTA2) and myeloid-lineage cells (CD68+, monocytes and macrophages). Myeloid cells were present in both eutopic endometrium and endometriotic lesions within stromal and epithelial compartments (Figure 1). ACTA2+ cells were present in the vasculature and stroma, although not in the immediate sub-epithelial stroma. Duplicate regions of interest were selected for each tissue and segmented based on fluorescence intensity for macrophages, epithelium, and stroma, resulting in 60 segments.

**Figure 1.**
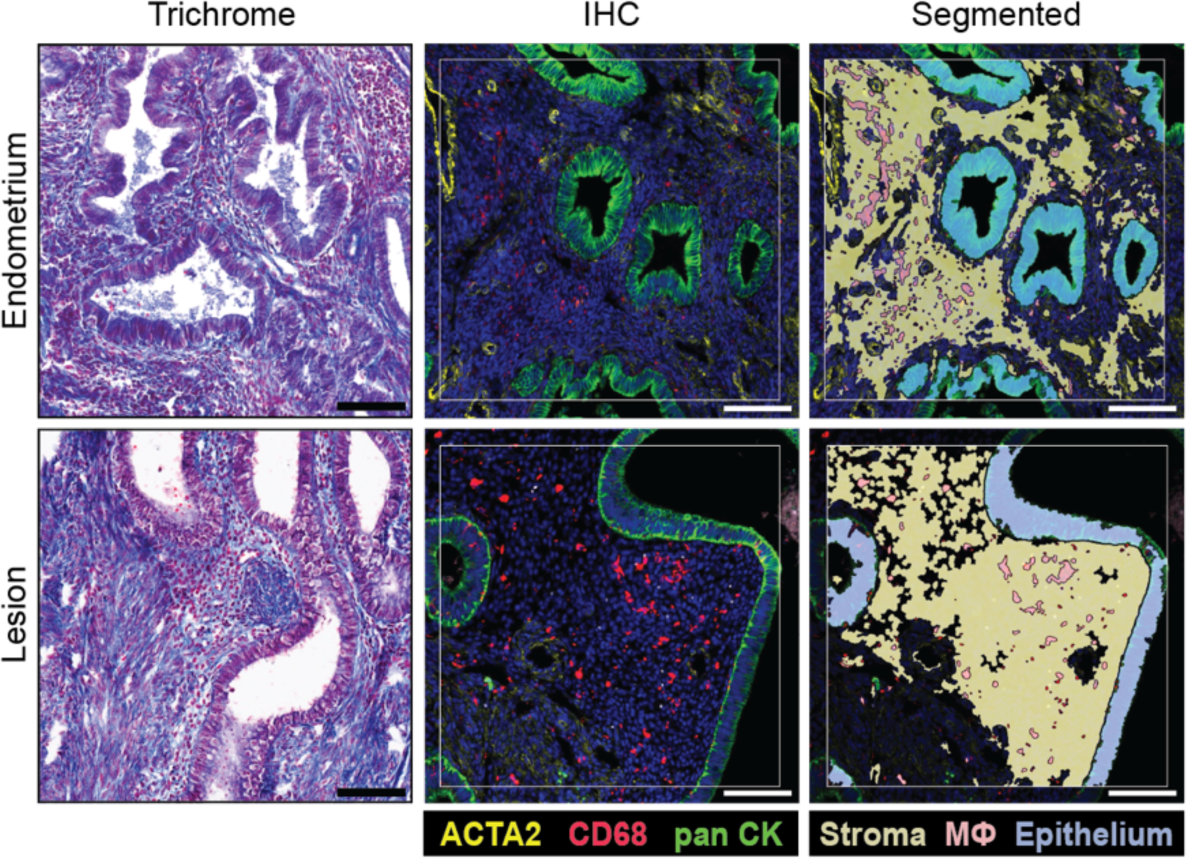
Segmentation strategy for spatial transcriptomics. Superficial peritoneal endometriotic lesions and matched endometrium (n = 10) were visualized with trichrome staining to confirm the presence of epithelial and stromal compartments. Tissues were immunostained for pan-cytokeratin, CD68, and ACTA2 for segmentation and spatial transcriptome collection on the GeoMx platform. Representative regions of interest (ROI) are shown for each tissue type. Duplicate ROI were collected for each tissue and stroma, epithelium, and macrophages were segmented from each for a total of 20 ROI and 60 segments. Scale bar = 100 µm.

**Table 1.**
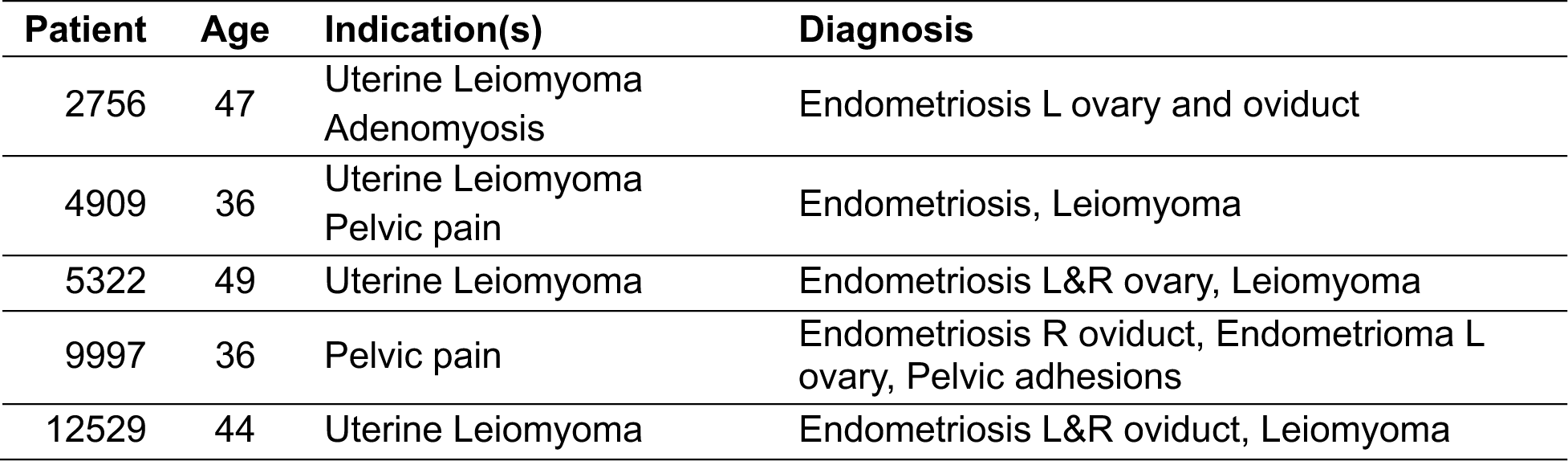
Patient demographics, indications for surgery, and post-operative diagnosis.

| Patient | Age | Indication(s) | Diagnosis |
| --- | --- | --- | --- |
| 2756 | 47 | Uterine Leiomyoma<br>Adenomyosis | Endometriosis L ovary and oviduct |
| 4909 | 36 | Uterine Leiomyoma<br>Pelvic pain | Endometriosis, Leiomyoma |
| 5322 | 49 | Uterine Leiomyoma | Endometriosis L&R ovary, Leiomyoma |
| 9997 | 36 | Pelvic pain | Endometriosis R oviduct, Endometrioma L ovary, Pelvic adhesions |
| 12529 | 44 | Uterine Leiomyoma | Endometriosis L&R oviduct, Leiomyoma |

Based on the limit of quantification, 10 segments were removed with a gene detection rate of lower than 10% (Figure 2A). Read counts were normalized for the remaining 50 segments using the quartile 3 (Q3) method. Separation between the geometric mean of the negative control probes, used to calculate the limit of detection, and Q3 of all gene read counts was confirmed at the segment and distribution levels to ensure compatibility with Q3 normalization (Figure 2B). Next, genes detected in less than 10% of segments were removed leaving 7,945 genes (Figure 2C). Overall gene expression across segments was visualized with a Uniform Manifold Approximation and Projection (UMAP) plot, which resulted in two major clusters (Figure 2D). Epithelial segments were contained in one cluster and the second cluster included both stroma and macrophage segments. Notably, segments from the same patient were found in close proximity within each cluster.

**Figure 2.**
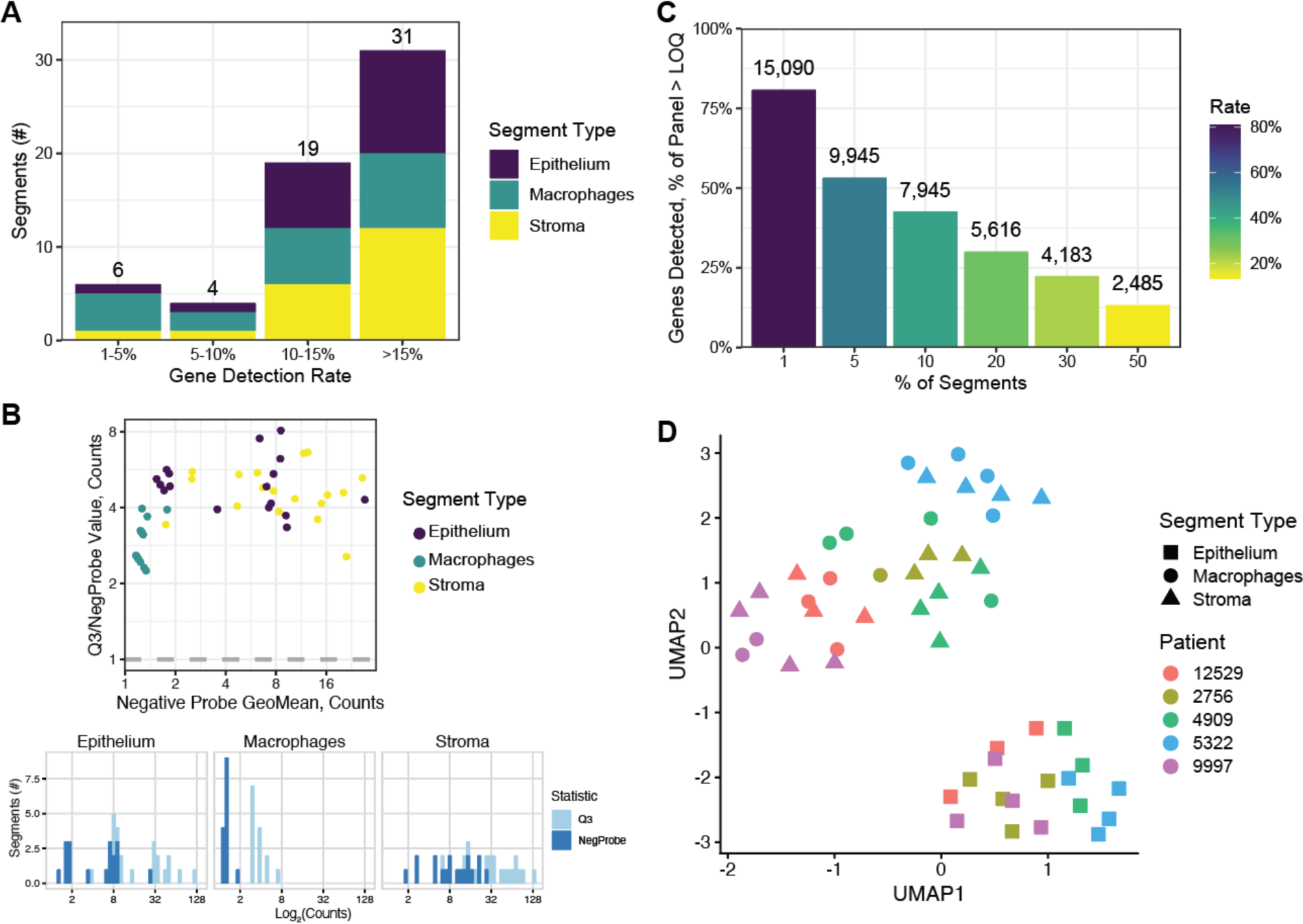
Spatial transcriptomics results from eutopic endometrium superficial peritoneal endometriotic lesions. (A) Bar plot of all 60 segments separated by gene detection rate. Segments with a detection rate of less than 10% were removed. (B) Confirmation of gene and negative probe count separation by segment in a scatter plot and by distribution for each cell type in histograms. Separation after gene and segment filtering is required for effective Q3 normalization. (C) Bar plot of detectable genes per segment used for filtering genes found in less than 10% of segments leaving 7,945 genes for further analysis. (D) Dimensionality reduction UMAP plot of all segments coded by shape and colored by patient. Epithelium clustered separately from stroma and macrophages.

### Epithelium and Stroma Marker Genes Are Similar in Lesions and Endometrium

We compared the epithelium and stroma segments by tissue type to confirm cell type separation and identify differences in the cell identity in the lesion tissue. The eutopic endometrial epithelium was compared to matching stroma and 1,372 DEG corresponding to cell type were found (Figure 3A). Similarly, there were 1,313 DEG in lesion epithelium versus stroma (Figure 3B) and 14 of the top 20 genes were in common. Gene expression patterns were similar in the endometrium and lesion tissues with a correlation coefficient of 0.81 (*p* < 2.2 × 10^-16^, Figure 3C). Stromal and epithelial biological process terms were distinct and consistent with cell type (Figure 4D). Vascularization, extracellular matrix, and cell adhesion terms were enriched in the stroma while epithelial cell differentiation and cell-cell junction terms were enriched in the epithelium. Terms were very similar in the stroma with 9 of 10 terms common between the eutopic endometrium and lesions. However, five terms were unique to the lesion epithelium, related to cell-cell adhesion and cell projections or cilia, and two terms to the eutopic endometrial epithelium, cellular transition metal ion homeostasis and circulatory system process.

**Figure 3.**
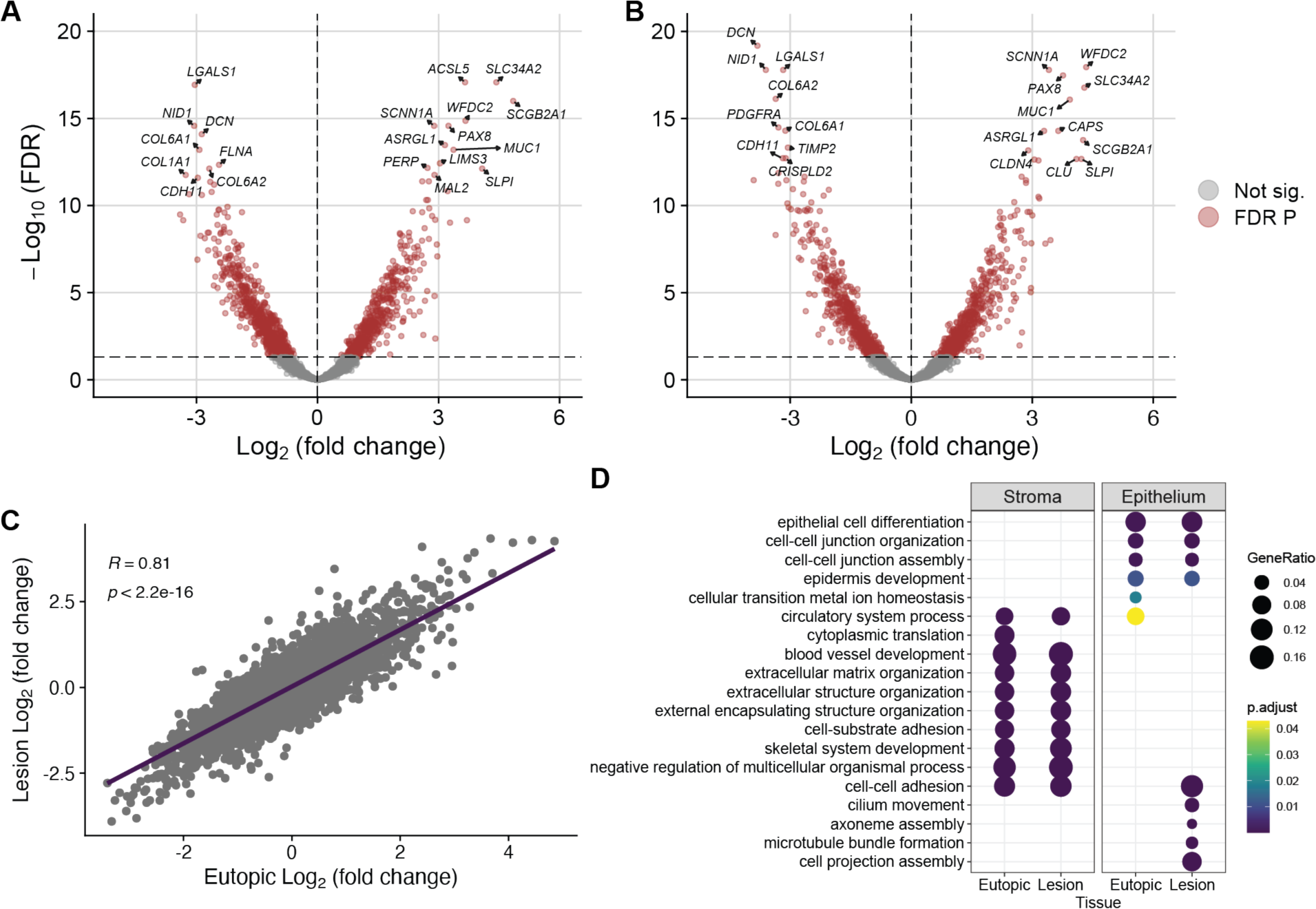
Epithelium versus stroma in eutopic endometrium and lesions. Volcano plots of differentially expressed genes (DEG) in the (A) endometrium and (B) lesions. The top DEG were similar in volcano plots with canonical epithelial genes like *MUC1*, *PAX8*, and *WFDC2* increased in both epithelia. (C) Genes enriched for each compartment were highly correlated in the eutopic endometrium and lesions. (D) Enriched gene ontology biological process terms were consistent with cell type, including extracellular matrix genes in the stroma and cell-cell junction genes in the epithelium, in the eutopic endometrium and lesions. Several terms related to cell-projections and cilia were unique, however, to the lesion epithelium.

**Figure 4.**
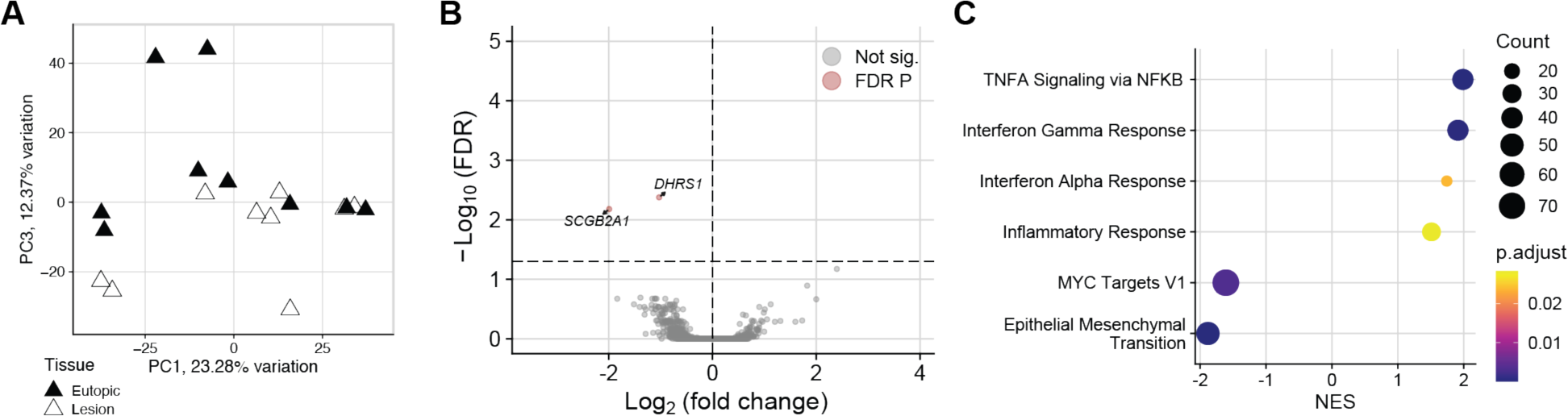
Endometriotic sub-epithelial stroma is minimally altered compared to eutopic endometrium. (A) Principal component (PC) plot of lesion stroma compared to eutopic endometrium showed group separation most clearly on PC3. (B) Volcano plot of two down-regulated genes in the endometriotic stroma. (C) Gene set enrichment analysis of hallmark pathways found four increased inflammatory gene sets and two decreased.

### Endometriotic Stroma is Minimally Altered

The sub-epithelial stroma from lesions was compared to matched eutopic endometrium to determine transcriptome alterations in endometriotic lesions. Separation of eutopic stroma from the lesions was strongest on PC3, accounting for 12.37% of variation in the dataset (Figure 4A). Two downregulated genes were identified in lesions (Figure 4B), *DHRS1*, dehydrogenase/reductase 1, and *SCGB2A1*, secretoglobin family 2A member 1, both related to steroid metabolism. Six hallmark gene sets [27] were significant in a GSEA, four increased and two decreased (Figure 4C). The increased gene sets were inflammation-related, TNFα signaling via NF-κB, interferon gamma response, interferon alpha response, and inflammatory response, and the decreased gene sets were MYC targets V1 and epithelial mesenchymal transition.

### Immune Response is Increased in the Endometriotic Lesion Epithelium

The endometriotic epithelium was compared to the matched eutopic glandular epithelium and was most separated from the eutopic endometrial epithelium on PC3, representing 9.45% of variation in gene expression (Figure 5A). Differential expression analysis found 76 DEG with 33 increased in the lesion epithelium and 43 decreased (Figure 5B). Notably, multiple immune-related genes were increased including, *C3*, or complement 3, *CD74*, *CLU*, or clusterin, and major histocompatibility complex class II members *HLA-DRB1* and *HLA-DPB1*. There were 10 significant hallmark gene sets from a GSEA with four terms increased and six decreased (Figure 5C). Among the increased gene sets were interferon gamma response and allograft rejection, which matched the increase in immune-related genes in the volcano plot. Genes decreased in response to damage by UV radiation, a source of tissue damage and inflammation, were decreased in lesions. Compared to the eutopic endometrium, glycolysis, angiogenesis, androgen response, and epithelial mesenchymal transition gene sets were decreased in the lesion epithelium. Genes increased in the lesion epithelium were enriched for 33 gene ontology (GO) biological process terms and nine of the top 10 were related to immune response (Figure 5D). In fact, nine of the 33 upregulated genes were associated with positive regulation of immune response: *CD74*, *HLA-DRB1*, *CLU*, *C3*, *HLA-DPB1*, *RPS19*, *HLA-DQB1*, *CFB*, and *VTCN1*.

**Figure 5.**
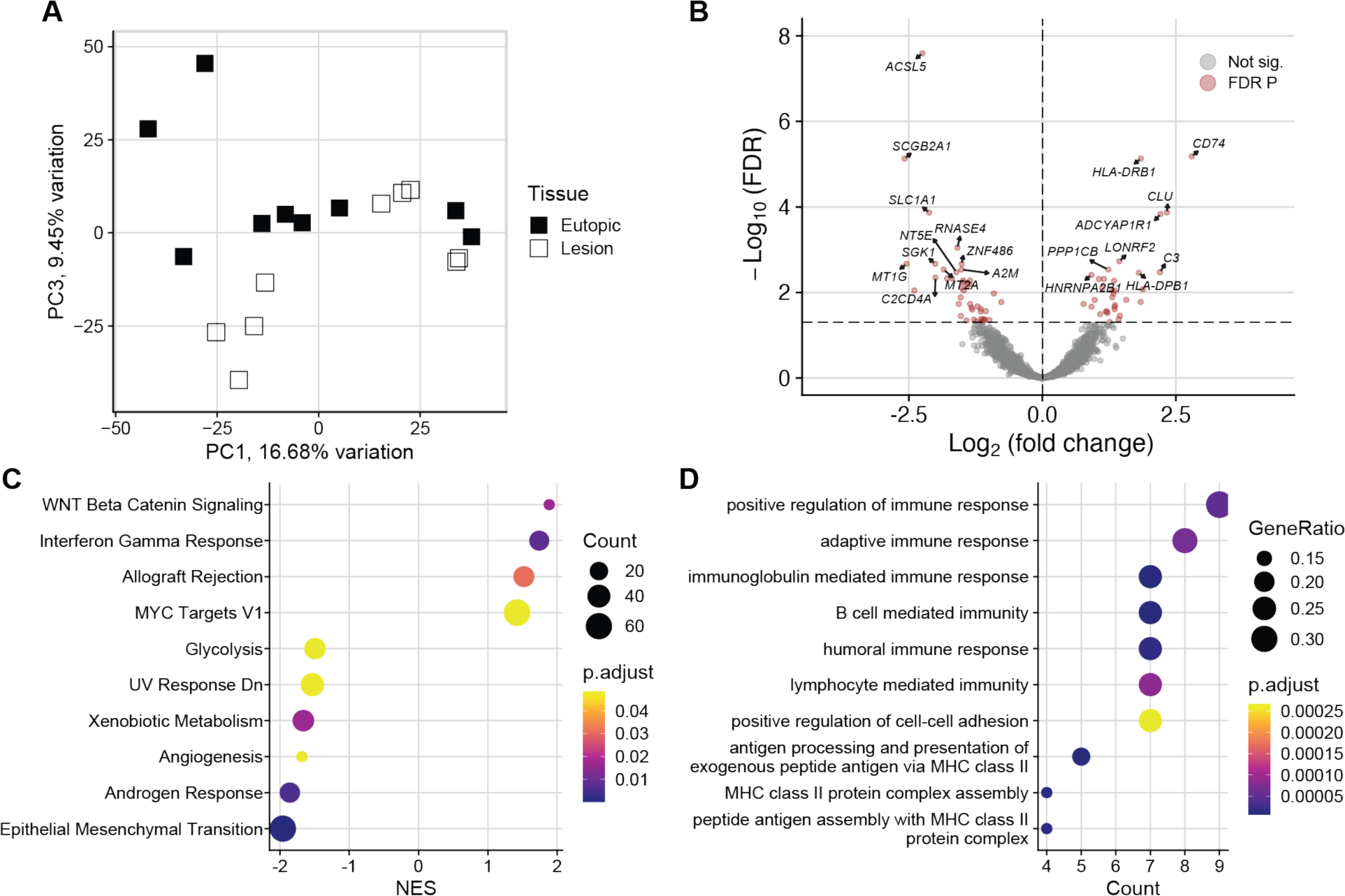
Immune response genes are increased in the endometriotic lesion epithelium. (A) Principal component (PC) plot of lesion epithelium compared to eutopic endometrium showed group separation most clearly on PC3. (B) Volcano plot of 76 differentially expressed genes (DEG) in the endometriotic epithelium. Of note, complement 3, *C3*, was one of the top increased DEG and several others were major histocompatibility complex class II related genes, like *CD74* and *HLA-DRB1*. (C) Gene set enrichment analysis of hallmark pathways found four increased gene sets and six decreased, indicating increased inflammation and proliferation. (D) Nine of the top 10 enriched gene ontology biological process terms were immune related.

### Neither Stroma nor Epithelium Clustered by Tissue

Hierarchical clustering dendrograms were plotted for the stromal and epithelial segments (Figures 6A-B) as an unsupervised measure of segment similarity. Neither cell type clustered by eutopic or lesion origin. Segments from two patients were clustered in the stroma and epithelium, that is lesion segments were more like eutopic tissue from the same patient than another lesion. Principal component analyses of the stroma and epithelium (Figures 6C-D) segments revealed that PC1 was driven by patient ID, representing 23.28% and 16.68% of variation in gene expression, respectively.

**Figure 6.**
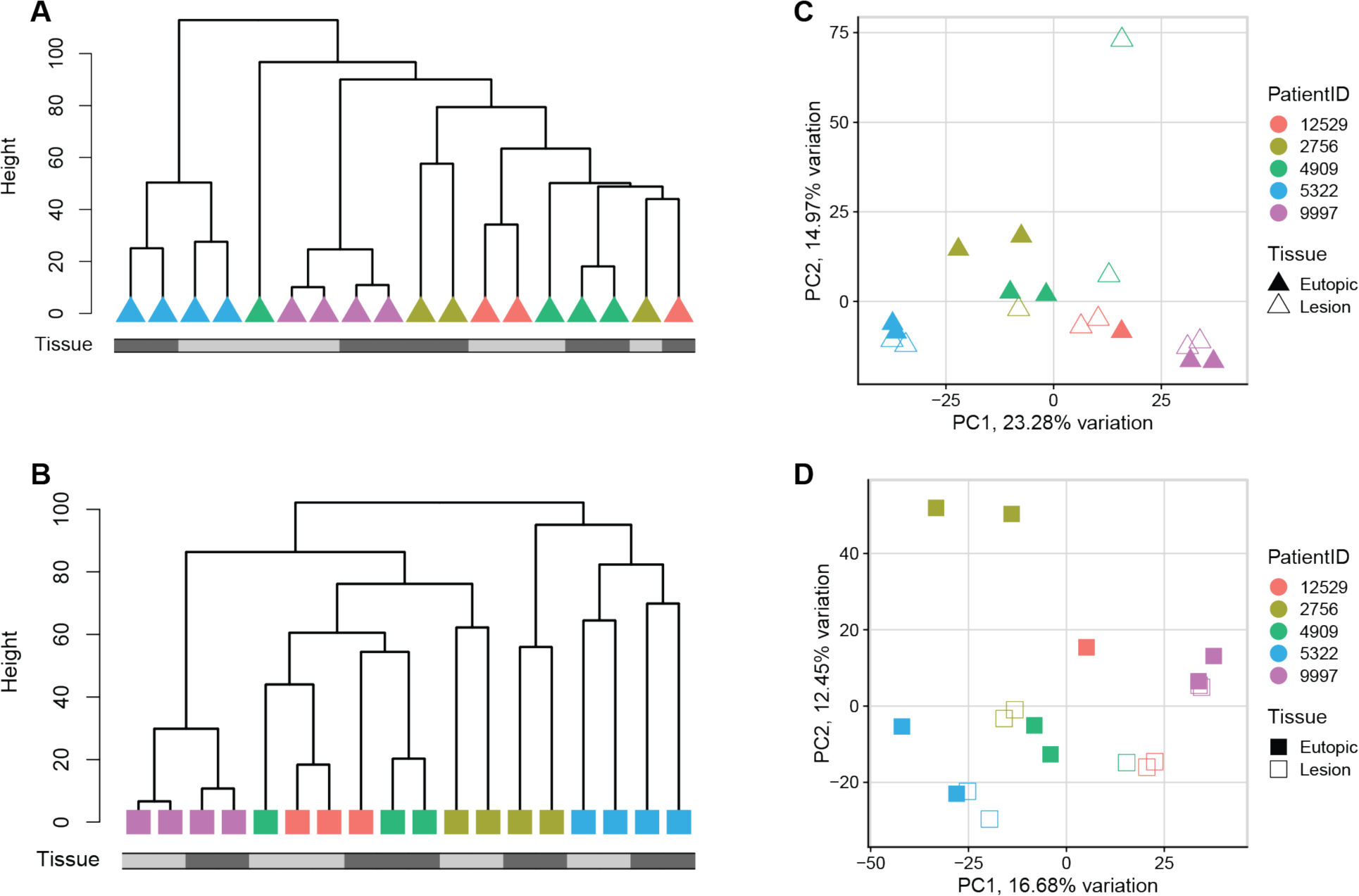
Neither stroma nor epithelium cluster by lesion versus eutopic endometrium. Hierarchical clustering dendrograms of (A) stroma and (B) epithelium segments found no clustering of segments by tissue of origin (endometrium = black bar). Principal component (PC) plots demonstrated that PC1, the largest component of variation, was correlated with patient identity.

### Macrophages are Altered in Peritoneal Lesions

Lesion macrophages were most separated from eutopic endometrium macrophages on PC3, representing 8.87% of variation in gene expression (Figure 7A). Differential expression analysis found 148 DEG with 35 increased and 113 decreased in the lesion macrophages (Figure 7B). Nine hallmark gene sets were significantly enriched from a GSEA with seven terms increased and two decreased (Figure 7C). The seven increased terms were associated with inflammation and included TNFα signaling via NF-κB, inflammatory response, allograft rejection, and IL2 STAT5 signaling. Gene sets that were decreased were myogenesis and epithelial mesenchymal transition. Neither macrophage number nor proximity to the epithelium was altered in lesions compared to eutopic endometrium (Supplemental Figure 1).

**Figure 7.**
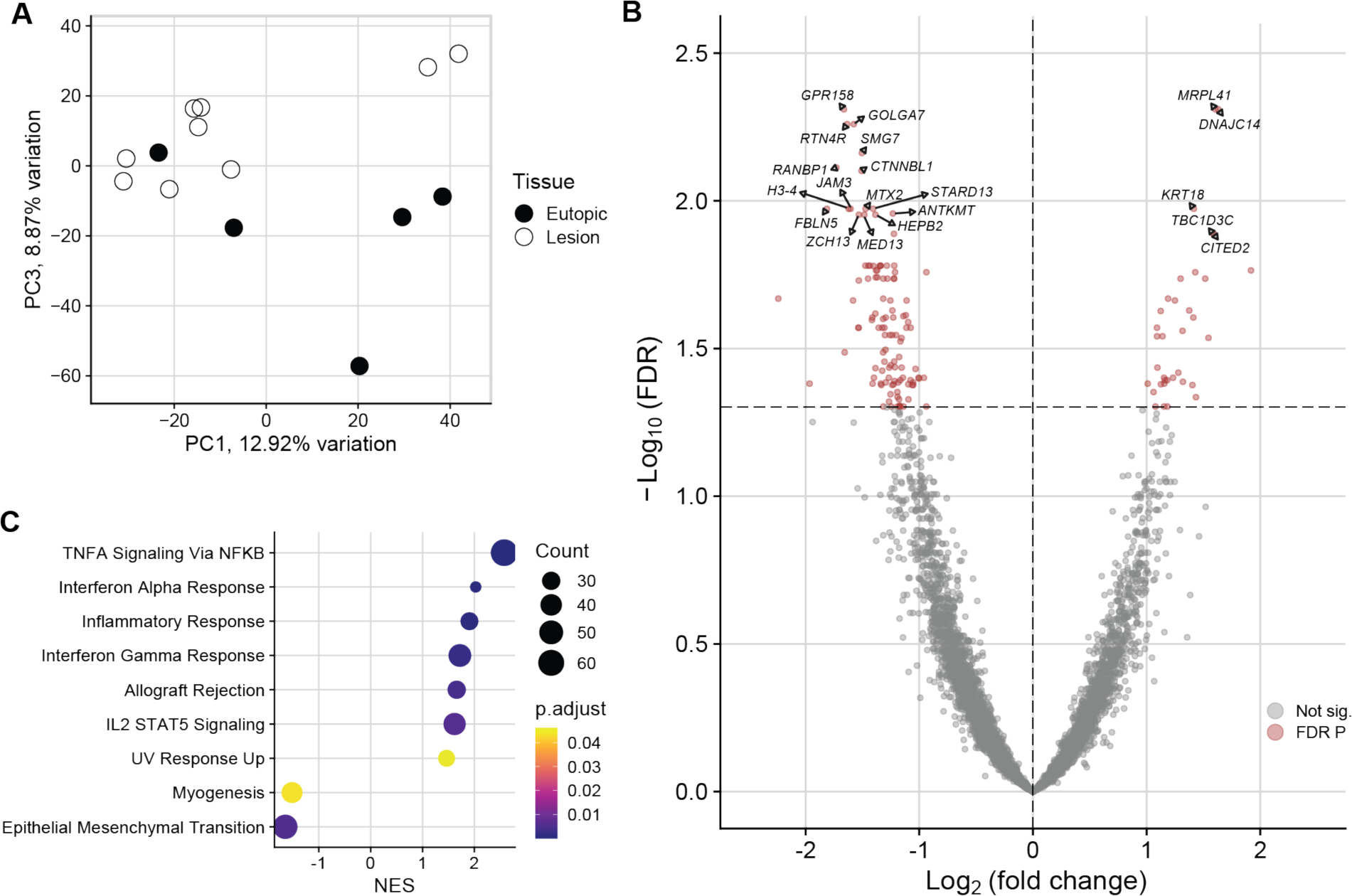
Macrophages are altered in endometriotic lesions. A) Principal component (PC) plot of lesion macrophages compared to eutopic endometrium showed group separation most clearly on PC3. (B) Volcano plot of 148 differentially expressed genes (DEG) in the endometriotic macrophages. (C) Gene set enrichment analysis of hallmark pathways found seven increased gene sets and two decreased, indicating increased inflammation likely reflective of the presence of monocyte-derived macrophages in the lesion tissue.

To provide context for our findings, we sought to compare our results with bulk sequencing data from human peritoneal lesions [21]. Human peritoneal lesions were separated from eutopic endometrium on principal component (PC) one (39.43% of variation) and 3,656 differentially expressed genes (DEG) were identified in an analysis of bulk RNA-sequencing data (Supplemental Figures 2A-C). Gene set enrichment analysis (GSEA) [28] found TNF⍺ signaling via NF-κB, myogenesis, immune response, and adipogenesis hallmark genes [27] increased, while estrogen response, glycolysis, and multiple cell cycle gene sets were decreased in endometriotic lesions (Supplemental Figure 1D).

### Ligand-Receptor Analysis Identified Epithelium-Macrophage Signaling Axis in Lesions

CellChat [29] ligand-receptor analysis was performed to determine the cell-cell communication landscape in endometriotic lesions and eutopic endometrium. Based on 976 significant ligand-receptor interactions, 29 signaling pathways were identified in total. Twenty-six pathways were present in lesions, 24 in the endometrium, and 21 pathways were common to both. Three pathways were unique to the endometrium, CXCL, SPP1, and TGF-β, and five unique to lesions, CD45, CD46, ITGB2, PTPRM, and SEMA3. A heatmap of signaling pathways consisting of all possible connections, inter- and intra-tissue, ranked by signaling strength (Figure 8A) found collagen signaling to be the strongest followed by MHC-II, laminin, macrophage migration inhibitory factor (MIF) and complement. The three top interacting cell types were lesion stroma, endometrium stroma, and lesion macrophages. Visualization of the cell-cell communication networks in the endometrium and lesions by weight in circle plots showed strong connections from the stroma compartment to the epithelium in both tissue types (Figure 8B). Based on combined network weights, confined to tissue type, the top senders were endometrial (0.172) and lesion stroma (0.170), and the top receivers were lesion macrophages (0.184) and epithelium (0.104). Particularly striking was a 3.70-fold increase in epithelium-to-macrophage signaling in lesions compared to eutopic endometrium. Interaction strength heatmaps (Figure 8C), confirmed the strong connection from the endometrial stroma received by the epithelium. In lesions, the strongest interaction was macrophage-macrophage signaling followed by epithelium-macrophage, surpassing stroma to epithelium signaling. These results point to a strengthened epithelium-macrophage signaling axis in endometriotic lesions.

**Figure 8.**
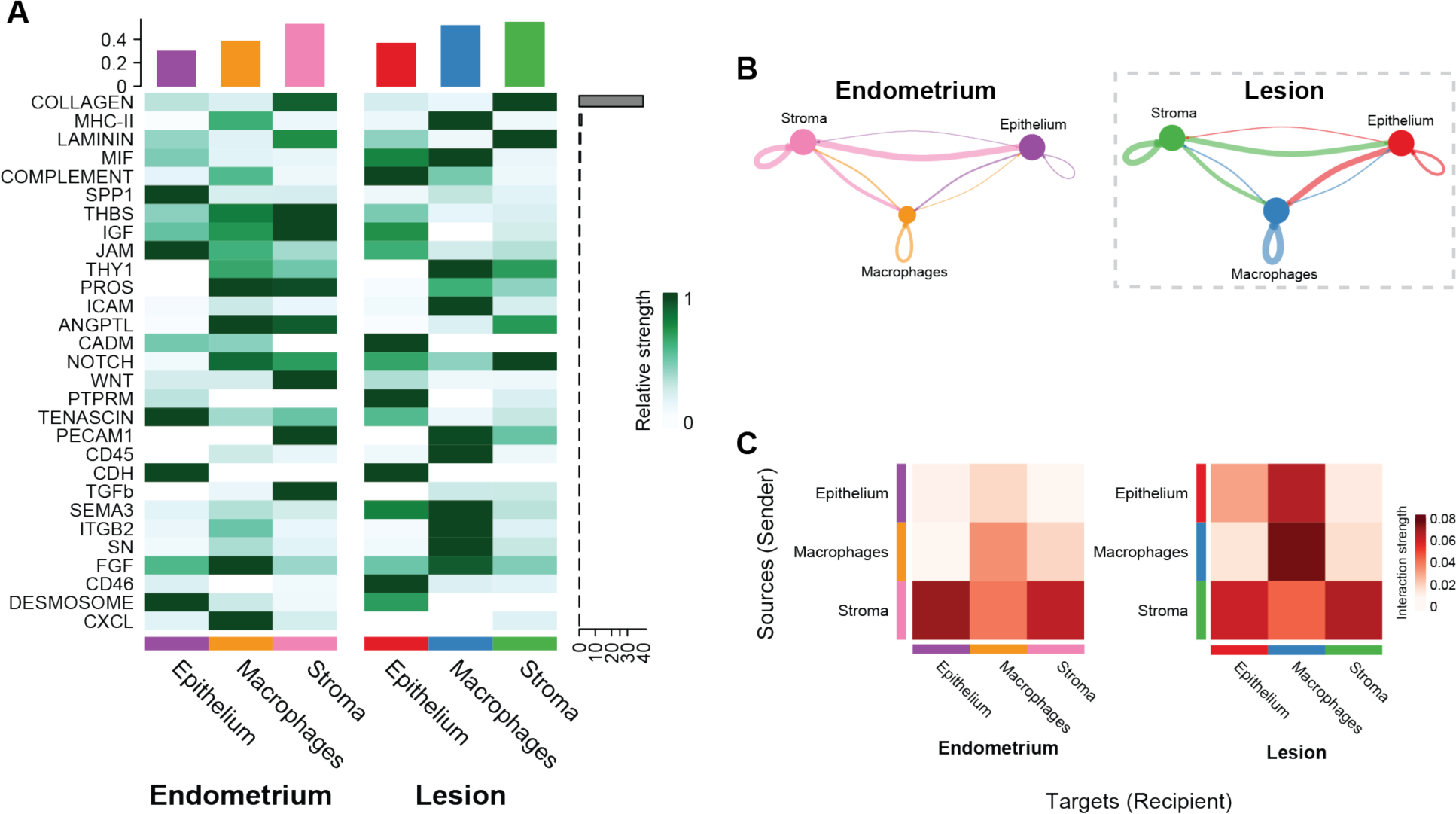
CellChat ligand-receptor analysis identified increased communication from the lesion epithelium to macrophages. (A) Signaling pathway heatmap of all 29 significant pathways identified by CellChat sorted by signaling strength, indicated by the gray bar. Interactions, combined incoming and outgoing, totaled for each cell type in the bar plot at the top of the heatmap and scaled per row. (B) Circle plots of communication networks including 24 pathways in the endometrium and 26 in lesions where line thickness corresponds to strength. Stroma to epithelium communication was strong in the eutopic endometrium and similarly present in the lesions. However, epithelium to macrophage communication was increased 3.7-fold in lesions. (C) Interaction strength heatmaps highlight the gain of epithelium to macrophage signals in lesions and strong macrophage-macrophage interactions.

Given that multiple inflammatory pathways were increased in lesions, including MHC-II and complement, and since increased signaling was noted between the lesion epithelium and macrophages, we sought to determine the source of inflammation-related ligands in lesions. A ligand-receptor bubble plot (Figure 9A) demonstrated that complement and MHC-II ligands were originating in the lesion epithelium and likely binding to receptors on macrophages. Within the stroma, MIF was the only visualized inflammatory pathway with likely communication probability indicating that inflammatory signals originated largely in the lesion epithelium. Indeed, a chord diagram of complement signaling (Figure 9B) highlighted the increased complement signaling present in lesions compared to the endometrium and the lesion epithelium as the source.

**Figure 9.**
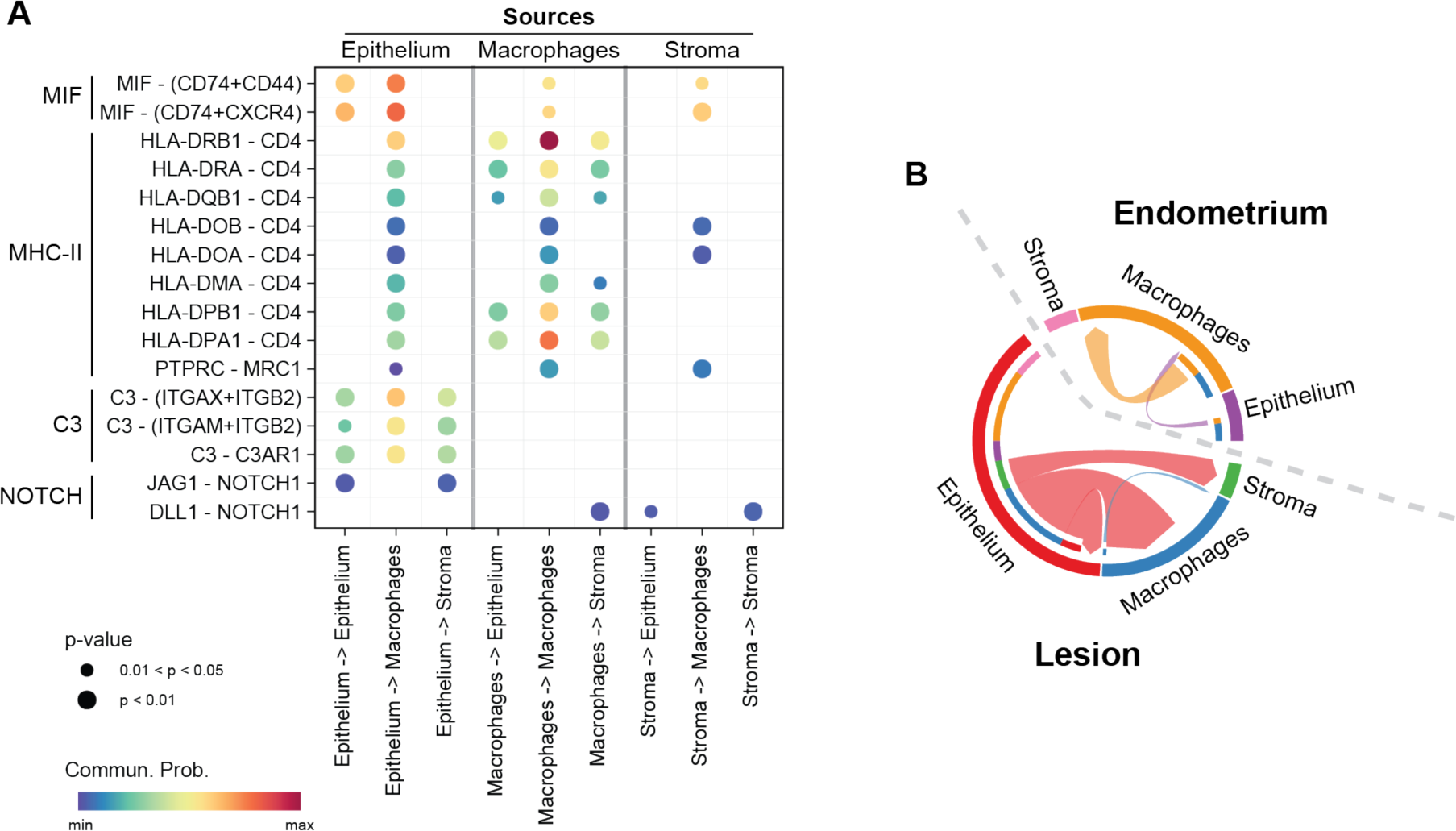
Inflammatory ligands originate in the endometriotic lesion epithelium. (A) Ligand – receptor dot plot of inflammatory pathways in endometriotic lesions. Pathway names are indicated on the left side of the plot followed by specific ligand and receptor interactions. All signals originating from the lesion epithelium are in the left three columns, macrophages in the center, and stroma on the right. Complement and major histocompatibility complex class II (MHC-II) pathways originated in the lesion epithelium and not the stroma. Macrophage migration inhibitory factor (MIF) signaling was strongest in the epithelium, but also sent from the stroma and macrophage cell types. (B) Complement signaling chord diagram of the weighted complement signaling pathway indicating increased signaling from the lesion epithelium compared to the eutopic endometrium epithelium.

## Discussion

Endometriosis is a complex reproductive disorder marked by the presence of endometrium-like tissue outside the uterus. Focusing on superficial peritoneal lesions, and excluding deep infiltrating or ovarian endometrioma disease, we hypothesized that endometrium-like tissue in these lesions would display an altered molecular phenotype influenced by the peritoneal microenvironment. To explore this hypothesis, we utilized spatial transcriptomic, receptor-ligand, and bulk RNA-sequencing analyses of endometriotic lesions compared to eutopic endometrium.

Bulk RNA-sequencing unveiled thousands of differentially expressed genes (DEGs) and numerous altered inflammatory pathways in lesions compared to the eutopic endometrium, highlighting the inflammatory nature of these lesions and the presence of fibrotic tissue. This observation supported the notion that endometrium-like tissue within the peritoneum would differ significantly from its uterine counterpart. Inflammatory cytokines are increased in the peritoneal fluid of women with endometriosis [30], likely influencing gene expression. Indeed, miRNAs, Notch signaling, aromatase enzymes, prostaglandins, and progestin responsiveness are altered in lesions compared to matched endometrial tissues [31–42].

Contrary to our initial expectations, spatial analysis revealed transcriptomic similarities between the three major cell types found in endometrium-like regions of lesions and patient-matched eutopic endometrium, with macrophages displaying the most substantial transcriptional difference. This lack of transcriptional divergence in the epithelium and stroma supports the idea that superficial peritoneal endometriotic lesions are derived from the endometrium following retrograde menstruation [8, 9]. Further, patient rather than tissue type, that is lesion or endometrium, was the primary determinant of gene expression based on principal component and UMAP plots. The differences observed in macrophage gene expression are likely due to the presence of peritoneal and monocyte-derived macrophages in lesions, compared to tissue resident macrophages in the eutopic endometrium [43–45].

The discrepancy between bulk RNA-seq and spatial analyses points to the inherent differences in methodologies. While bulk RNA-seq provides a comprehensive view of gene expression changes across the entire homogenized tissue, spatial transcriptomics offers a targeted approach focusing on localized gene expression patterns within distinct regions of interest, such as the endometrium-like structures of lesions. This targeted approach resulted in comparatively few DEG, indicating that the high difference in transcriptomic profiles between lesions and eutopic endometrium found in bulk RNA-seq may not be attributable to endometrium-like tissue in lesions but from the peripheral tissue. This highlights the importance of spatially resolved data in heterogenous tissues like endometriotic lesions. For example, we expect many of the DEG identified in bulk RNA-seq originate from the fibrotic regions of lesions and reflect sustained inflammation and the presence of activated fibroblasts [9, 16]. Interestingly, our findings suggest that the sub-epithelial stroma in superficial peritoneal lesions remains largely unaltered and could serve as an internal control for future spatial experiments, bypassing the requirement for matched eutopic stromal tissue. The paired design of this study increased power with a limited sample size. However, the lack of healthy control endometrium and unknown time since symptom onset are notable caveats and may limit the generalizability of the findings, particularly for the initiation of disease. As such, further research is necessary to elucidate the molecular mechanisms underlying the early pathogenesis of superficial peritoneal endometriosis.

The spatial transcriptomic technique, sequencing hybridized probes from specific areas of FFPE tissue, resulted in an overall reduced transcriptome size after removing genes with low counts across segments. The spatial transcriptome included 7,945 genes while the bulk RNA-sequencing analysis included 17,405 genes. Consequently, many of the lowest expression genes from the bulk RNA-sequencing data were filtered from the spatial transcriptome before DEG analysis. The advantage of spatial transcriptomics, however, is gaining information from genes expressed only by a subset of cells or a single cell type that would be lost in the bulk tissue. This is evidenced by the inclusion of 551 genes in the spatial transcriptome that were considered as not expressed in the bulk RNA-sequencing analysis. Further, spatial transcriptomics indicated increased proliferation of the lesion epithelium while bulk RNA-sequencing found an overall reduction in cell cycle gene sets. The epithelium of lesions is likely to be a minor contributor to gene expression from homogenized tissue since lesions often have little or even no remaining epithelial component. Interestingly, these results suggest the lesion epithelium may proliferate in response to inflammatory peritoneal environment while stroma reduces proliferation and may even become senescent [46] . A growing body of evidence has found that senescence plays a role in multiple pathologies and may contribute to endometriotic lesions [47–49].

Receptor-ligand analysis identified an almost four-fold increase in signaling between epithelium and macrophages in lesions compared to the endometrium, including multiple inflammatory pathways. MHC-II ligands expressed in the lesion epithelium were predicted to communicate to macrophages through the CD4 receptor. Notably, human monocytes and macrophages express CD4, unlike the mouse, serving as an important target for HIV-1 infection [50–52]. The canonical macrophage marker from this study, CD68, is expressed by monocytes and macrophages [53]. Although monocytes rapidly differentiate into macrophages within a tissue, some monocytes were likely included in the spatial transcriptome analysis. Canonically, MHC-II binding of the CD4 receptor on monocytes promotes differentiation into macrophages [51], thus increased MHC-II in the lesion epithelium would drive macrophage differentiation from circulating monocytes homing to areas of hypoxia and inflammation. Increased expression of MHC class II expression in the epithelium aligns with recent single cell RNA-sequencing analysis of ovarian endometriosis, which found increased MHC class II expression in the epithelium [54].

Macrophage migration inhibitory factor (MIF) is constitutively expressed by many cell types and has pleiotropic autocrine and paracrine proinflammatory effects on tissues [55]. MIF signaling in macrophages sustains proinflammatory function by inhibiting apoptosis, counteracting the immunosuppressive effects of glucocorticoids, and promoting secretion of prostaglandin E2 [55, 56]. Additionally, MIF is released by monocytes, macrophages, and other immune cells upon exposure to proinflammatory mediators, such as C3, to reinforce a local proinflammatory environment [57–59].

*Complement 3* expression was increased in bulk RNA-sequencing results (3.7-fold, FDR *p* = 0.0003) and found in the lesion epithelium from spatial transcriptomic analysis (4.6-fold, FDR *p* = 0.003). Endometriotic lesions were originally reported to express Complement 3 (C3) by Weed and Arquembourg in 1980 [60] and later found to be secreted by the lesion epithelium [61, 62]. In fact, serum levels of C3 are elevated in patients with endometriosis and C3 was proposed as a potential diagnostic biomarker [63]. The complement system is a signaling cascade that plays a pleiotropic role in immune response, including production of proinflammatory mediators and recognition and clearance of pathogens by lysis and enhanced phagocytic uptake by immune cells [64]. The cascade of over 50 proteins is characterized by a series of protein cleavage events that produce inflammatory cytokines and the components for subsequent steps. Multiple stimuli can activate the complement system, but all pathways converge on activation of C3 by a convertase. Although the presence of C3 in endometriotic lesions has been known for 40 years, its potential contribution to the disease is unclear [65]. Following cleavage, the resulting fragments are referred to by appending the letter “a”, for the smaller fragment, or “b”, for the larger fragment, to the protein name, for example, C3a.

C3a and C3b are produced following activation of C3 by a C3 convertase, itself the product of prior cleavage events. C3b induces phagocytic activity of macrophages, but lesion tissue is refractory to complement targeting through multiple mechanisms. CD47, otherwise known as the “don’t eat me signal”, is expressed by tissue and binds to the SIRPɑ receptor on macrophages, inhibiting phagocytosis [66, 67]. Complement regulatory proteins, such as CD35, CD46, and CD55, inhibit complement attack by accelerating breakdown or direct cleavage of complement components critical for continuation of the cascade [64]. In the event of complement activation, CD59, another complement regulatory protein, inhibits formation of the membrane attack complex and prevents cell lysis [68]. Thus, multiple mechanisms exist within the lesion to prevent direct complement attack but complement signaling promotes a localized proinflammatory environment. The proinflammatory cytokine C3a, released following C3 cleavage, can activate a pro-repair or tumor-associated macrophage (TAM) phenotype by binding to C3AR1, as found in the ligand-receptor analysis [69]. This provides a mechanism by which C3 activation from the superficial peritoneal endometriotic lesion epithelium promotes the pro-lesion macrophage phenotype described as pro-repair, tumor-associated, or M2. These macrophages then contribute to lesion tissue remodeling including neovascularization and fibrosis [18, 70, 71]. Additionally, semaphorin 3B was found in the lesion epithelium signaling to macrophages and could further promote the pro-disease TAM phenotype since SEMA3B increased expression of pro-repair macrophage markers in a model of rheumatoid arthritis [72].

Activated complement signaling in tissue induces a diffuse inflammatory state that includes increased prostaglandins, also upregulated by MIF [65]. Increased prostaglandins can activate steroidogenic enzymes and, potentially the production of estradiol and estrone by the lesion [31, 33, 35, 73]. Finally, increased estrogens can activate complement in lesion tissue [74] resulting a positive feedback loop, common in pathogenic signaling. In this manner, the complement signaling is sustained by local estrogenic activity and could sustain local inflammation and the presence of pro-repair macrophages until such time as the lesion becomes completely fibrotic with a tissue stiffness that no longer supports cell growth and the lesion becomes scar tissue and contributes to peritoneal adhesions. In support of complement signaling promoting lesion pathogenesis, C3 null uterine tissue formed fewer lesions after 21 days in a mouse model of endometriosis [75].

The comparison of lesions to eutopic endometrium yielded intriguing insights into the molecular and cellular dynamics underlying lesion pathophysiology. Firstly, despite previous findings from bulk tissue analysis suggesting significant differences between endometrium-like tissue in lesions and eutopic endometrium, our spatial transcriptomic data reveal a surprising similarity between these two tissues. The lack of drastic differences may be attributed to the composition of the excised lesions used in bulk RNA-sequencing, with endometrium-like tissue comprising only a small proportion of the lesion, such that differential expression was largely attributable to the peripheral stroma and fibrotic tissue. Notably, this difference highlights the importance of employing advanced techniques, like spatial transcriptomics, to gain an improved understanding of lesion biology.

Secondly, spatial transcriptomics has provided novel insights into the intricate cellular crosstalk within lesions. Our data indicate signaling between the stroma and epithelium, as well as between the lesion epithelium and macrophages. Notably, the epithelium emerged as a central player in driving inflammation within the lesion, promoting inflammatory signaling independent of the stromal compartment. Moreover, the presence of CD68+ cells amid epithelial cells highlights the physiological relevance of cell-cell interactions across different cell types.

Drawing from these observations, we present a model wherein the superficial lesion epithelium orchestrates inflammatory signaling within the lesion while promoting a pro-repair phenotype in infiltrating macrophages. Spatial transcriptomics facilitated the identification of inflammatory signaling emanating from the epithelium, including C3, MIF, and MHC class II. This novel insight sheds light on the pathological consequence of C3 expressed by the epithelium in modulating macrophage phenotype to support lesion development. In summary, our study underscores the critical role of considering spatial context and cellular interactions within lesions to unravel the complexities of superficial peritoneal endometriotic lesions. Further investigation is imperative to elucidate the specific mechanisms governing epithelium-driven inflammation and its implications for disease progression and therapeutic intervention.

## Methods

### Human Tissues

All protocols involving tissue collection were approved by the institutional review board (IRB) Committee of Ponce Health Sciences University, School of Medicine (PHSU). Endometriotic lesions and matched eutopic uterine tissues from 10 patients were obtained as archived deidentified formalin-fixed paraffin-embedded (FFPE) tissue blocks [76–78]. All tissues were previously evaluated by a pathologist to confirm the diagnosis of endometriosis and menstrual cycle stage of eutopic endometrium using Noyes criteria [79].

### Histology

Tissue blocks were sectioned and trichrome stained (StatLab #KTTRBPT) to confirm the presence of defined epithelial and stromal compartments in lesions. Blocks were serial sectioned at 6 µm thickness and floated onto positively charged microscope slides. Tissues were attached to the slides prior to processing by heating on a slide warmer for 15 minutes. Tissues were then deparaffinized and rehydrated through three changes each of xylenes for five minutes and 100% ethanol for one minute before a rinsing in running tap water for one minute. Slides were incubated in Bouin’s fluid at room temperature overnight. Slides were rinsed in running tap water for three minutes and immersed in modified Mayer’s hematoxylin stain for four minutes at room temperature. After a three-minute rinse in running tap water, slides were immersed in one step trichrome stain for five minutes and rinsed in running tap water for five seconds. Slides were then dehydrated through three changes of 100% ethanol for one minute, cleared in three changes of xylenes for one minute, and a coverslip was applied with Permount mounting medium (Fisher Scientific #SP15).

### Spatial Transcriptomics

Human tissue slide preparation, library generation, and sequencing for digital spatial profiling (DSP) on the NanoString GeoMx DSP instrument were performed by the Van Andel Institute Histology and Genomics Cores. Sections were cut at 5 μm thickness and mounted on plus-charged slides (Epredia Colormark Plus #CM-4951WPLUS-001). Slides were baked at 60°C for one hour and stored at 4°C in a vacuum-sealed container containing desiccant for up to two weeks. All subsequent steps were performed under RNase-free conditions with DEPC-treated water. Slides were deparaffinized with three sequential five-minute washes in xylenes, followed by two washes in 100% ethanol for five minutes, one wash in 95% ethanol, and one wash in 1X PBS. Antigen retrieval was performed in target retrieval reagent (EDTA, pH 9.0; Invitrogen #00-4956-58) diluted to 1X in the BioGenex EZ-Retriever System for 10 minutes at 95°C. Slides were then washed with 1X PBS for five minutes. Slides were then incubated in 1 μg/mL proteinase K 10 minutes at 37°C and washed in 1X PBS for five minutes at room temperature. Slides were fixed for five minutes in 10% neutral buffered formalin followed by two washes in NBF stop buffer, for five minutes each, and one wash in 1X PBS for five minutes. Slides were then incubated with UV-photocleavable Human Whole Transcriptome Atlas hybridization probes diluted in buffer R (GeoMx RNA Slide Prep FFPE-PCLN kit, #121300313) in a hybridization oven at 37°C for 16-20 hours. Following probe incubation, slides were washed with stringent buffer (1:1, formamide:4X SSC buffer; Thermo Fisher #AM9342; Sigma-Aldrich #S6639) at 37°C twice for 25 minutes each. Then slides were washed twice in 2X SSC buffer. Slides were blocked in 200 μL buffer W (GeoMx RNA Slide Prep FFPE-PCLN kit) for 30 minutes and incubated at 4°C overnight with SYTO 13 nucleic acid stain (500 nM; Thermo Fisher Scientific #S7575) and antibodies for pan-cytokeratin (panCK; 1 µg/mL; Novus Biologicals #NBP2-33200AF488), smooth muscle actin (ACTA2; 1.25 µg/mL; Abcam #ab202368), and CD68 (0.5 µg/mL; Santa Cruz Biotechnology #sc-20060AF647) diluted in buffer W. Slides were washed four times in 2X SSC buffer for 3 minutes each and placed in the NanoString GeoMx DSP instrument.

Duplicate 500 µm^2^ regions of interest (ROI) were selected for each based on fluorescent imaging on the GeoMx DSP instrument (NanoString Technologies). ROI were drawn in the central portion of the endometrium stratum functionalis, to include glandular and not luminal epithelium, for eutopic uterine tissue and in lesions areas with defined epithelium. ROIs were then segmented in order of macrophages (CD68+), epithelium (panCK+), and stroma (ACTA2-) areas of illumination (AOI) for individual collection. AOI were processed sequentially on the instrument using UV light focused through each AOI that released the photocleavable oligos into wells of a 96 well plate. After AOI collection, indexing primers were hybridized during NanoString library preparation. Quality and quantity of the finished library pools were assessed using a combination of Agilent DNA High Sensitivity chips (Agilent Technologies, Inc.) and the QuantiFluor dsDNA System (Promega Corp.). Paired-end, 50 bp sequencing was performed on an Illumina Novaseq 6000 sequencer using an S2, 100 cycle sequencing kit v1.5 (Illumina Inc.). Base calling was done by Illumina RTA3 and output was demultiplexed and converted to fastq format with bcl2fastq v1.9.0.

### Spatial Data Analysis

Fastq files were converted to digital count conversion (DCC) files using GeoMx NGS Pipeline v2.3.3.10. Sample annotation, PKC configuration, and DCC files were imported with GeomxTools (v3.2.0) in R to generate a NanoStringGeoMxSet object. Read counts with a value of zero were shifted to one for downstream transformations and quality control (QC) analyses. Each AOI, or segment, was assessed for sequencing and tissue quality using the parameters listed in **Box 1**. Outlier probes were then excluded before counts were collapsed to gene-level data (18,677 genes) by geometric mean. The limit of quantification (LOQ) was determined per segment based on the distribution of negative control probes and set at two geometric standard deviations above the geometric mean with a minimum value of two. Segments with a gene detection rate less than 10% were filtered before removing genes detected in less than 10% of segments. Separation between the upper quartile 3 (Q3) of gene counts and the geometric mean of the negative probes was confirmed before count normalization using the quartile 3 (Q3) method.

Normalized read counts were log_2_-transformed and dimension reduction was performed by uniform manifold approximation and projection (UMAP) [80] with the umap package (v0.2.10) with the random state set to 42. Differential expression analyses were completed using log_2_-transformed normalized read counts with a linear model approach in the limma package (v3.54.2) [81]. A linear model was fit for each gene with lmFit before an empirical Bayes test was completed using patient as a covariate. P-values were corrected for multiple testing with the Benjamini Hochberg method included in the decideTests function of limma. Genes were considered differentially expressed when the adjusted p-value was below 0.05. Principal component analysis, volcano plots, gene set enrichment analysis, and over-representation tests were completed as described for bulk RNA-sequencing data.

### Macrophage Quantification

The macrophage count distance between from epithelium was manually measured for each segment using the line tool in ImageJ software [82]. The global scale was set by measuring the scale bar on the image (1 pixel = 2.51 µm). For each image, the green epithelium and red macrophage channels were merged for analysis. A line was drawn from the center of each macrophage to the center of the closest epithelial cell and the measurements. Based on these measurements the macrophages count and the average distance for each segment was calculated. Data were analyzed in Prism (v10.2.2, GraphPad Software LLC) following confirmation of normality (Shapiro-Wilk, p > 0.1). To determine if there was a relative increase in lesions compared to the matched endometrium, macrophage count was tested using a ratio paired t-test. Macrophage distance to the nearest epithelium was tested using a paired t-test.

### Bulk RNA-sequencing Analysis

Read counts per gene were downloaded from eutopic endometrium (n = 7) and peritoneal endometriotic lesion samples (n = 6) for series GSE179640 [21] from the NCBI Gene Expression Omnibus repository. Files were parsed with R (version 4.2.0) and all subsequent analyses, unless otherwise noted, were completed in R using RStudio [83]. Differential expression analysis was conducted using the edgeR-robust method (v4.0.3) [84, 85]. Genes with low counts per million (CPM) were removed using the filterByExpr function from edgeR. Multidimensional scaling plots, generated with the plotMDS function of edgeR, were used to investigate group separation prior to statistical analysis. Principal component analysis of normalized log_2_-transformed CPM was conducted with the PCAtools package (v2.10). The biplot and pairsplot functions were used to visualize sample separation across principal components one through five. Differentially expressed genes (DEG) were identified based on Benjamini-Hochberg false discovery rate (FDR) p-values less than 0.05. The hclust function was used for hierarchical clustering (ward.D2) of Euclidean distances and plotted with the dendextend package (v1.17.1). A volcano plot was generated with EnhancedVolcano (v1.16) and the top 20 DEG, based on FDR p-value, were labeled. Gene set enrichment and over-representation analyses were completed with clusterProfiler package (version 4.6.0) [86] using mSigDB [27] reference databases.

### Receptor-Ligand Analysis

Quantile-normalized read counts were used for receptor-ligand analysis with the CellChat package (1.6.10) in R [29]. Significant genes and interactions were identified using the functions identifyOverExpressedGenes and identifyOverExpressedInteractions. Communication probabilities were calculated between cell types with raw input counts option enabled, as recommended for bulk-sequencing data. The resulting interactions were visualized using the included plotting functions. Weighted communication probabilities were subset for eutopic endometrium or lesions before calculating tissue-specific communication interaction strengths.

### Data Availability

Sequence data from the spatial transcriptomics experiment were deposited in the NCBI Gene Expression Omnibus and available under series GSE263897.

## Acknowledgements

The authors would like to sincerely thanks all members of the Fazleabas laboratory and the Van Andel Genomics core, particularly Becca Siwicki, for their contributions to this project. This work was supported by NanoString Technologies, Incorporated (GWB) and National Institutes of Health grants F32HD104478 (GWB), R01HD099090 (ATF), R01HD050559 (IF), U56CA126379 (IF), and T32HD087166 (GWB, ATF). The content is solely the responsibility of the authors and does not necessarily represent the official views of the National Institutes of Health.

## Author contributions

Conceptualization, G.W.B., E.G., and A.T.F.; Data curation, G.W.B, Z.F., Z.B.M, and I.F.; Formal analysis, G.W.B, Z.F., E.L.V, and Z.B.M.; Funding acquisition, G.W.B, I.F, and A.T.F.; Investigation, G.W.B and E.L.V.; Methodology, G.W.B, Z.F., E.L.V, and Z.B.M; Resources, I.F. and A.T.F; Software, G.W.B, Z.F., and Z.B.M; Supervision, E.G. and A.T.F; Visualization, G.W.B; Writing – original draft, G.W.B and A.T.F.; Writing – review & editing, G.W.B, Z.F., E.G., I.F., and A.T.F.

## Declaration of generative AI and AI-assisted technologies in the writing process

During the preparation of this work, the authors used ChatGPT (v3.5) for proofreading and to improve readability. After using this service, the authors reviewed and edited the content as needed and take full responsibility for the content of the publication.

---

**Box 1.**
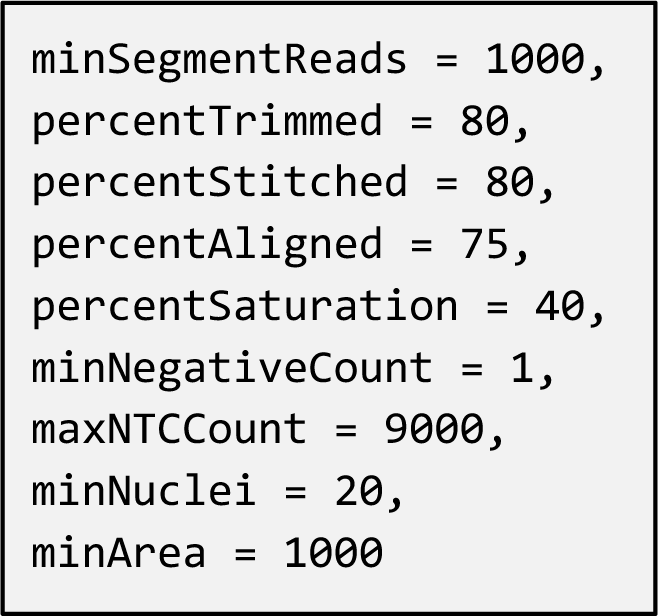
Spatial transcriptomics quality control parameters for segment filtering.

**Supplemental Figure 1.**
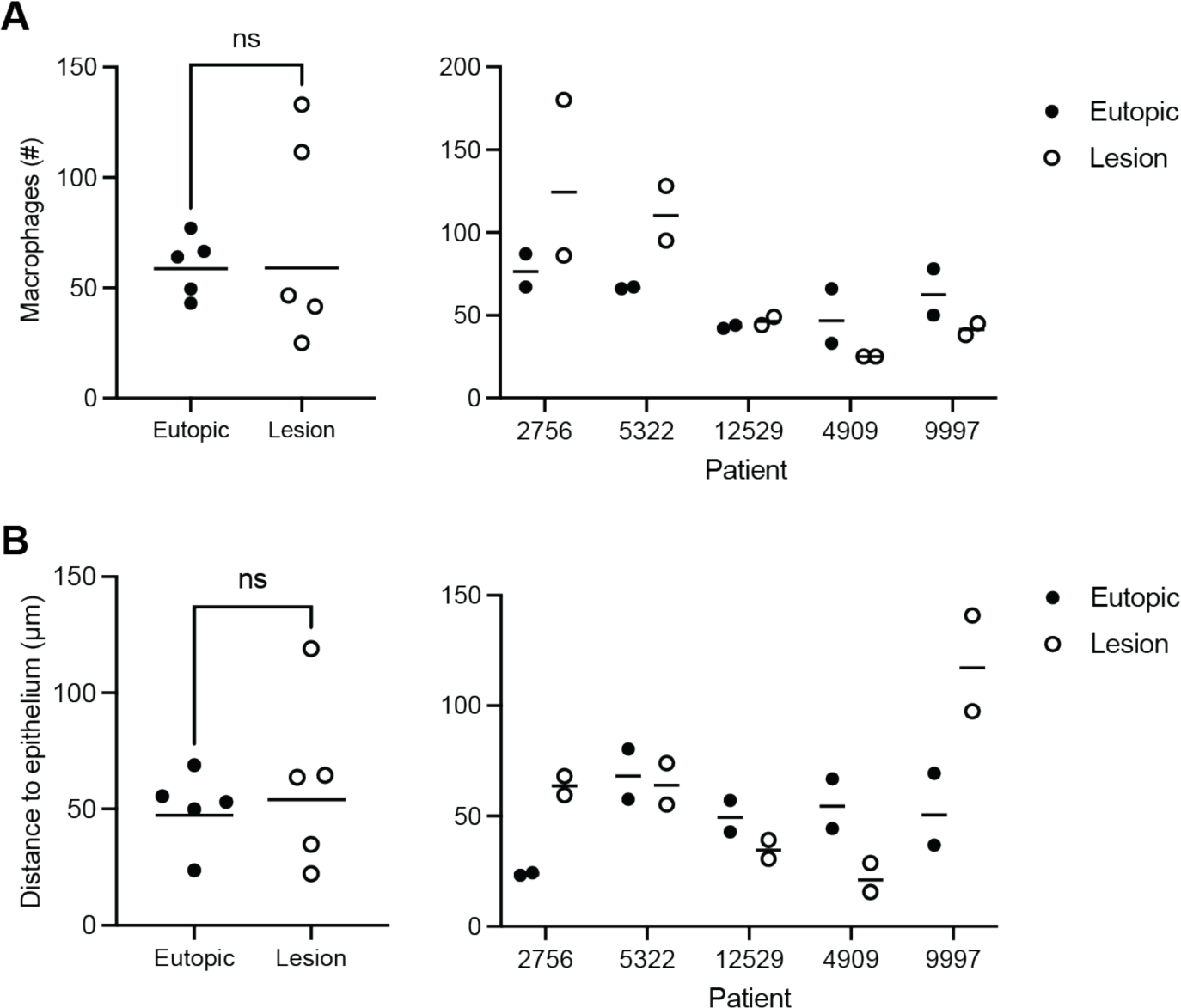
Quantification of macrophages in spatial transcriptomic segments from eutopic endometrium and superficial peritoneal lesions. Data are shown by group and individual patient. (A) Macrophage number was not increased in lesions (p = 0.98) and (B) did not appear to be located closer to the lesion epithelium (p = 0.59).

**Supplemental Figure 2.**
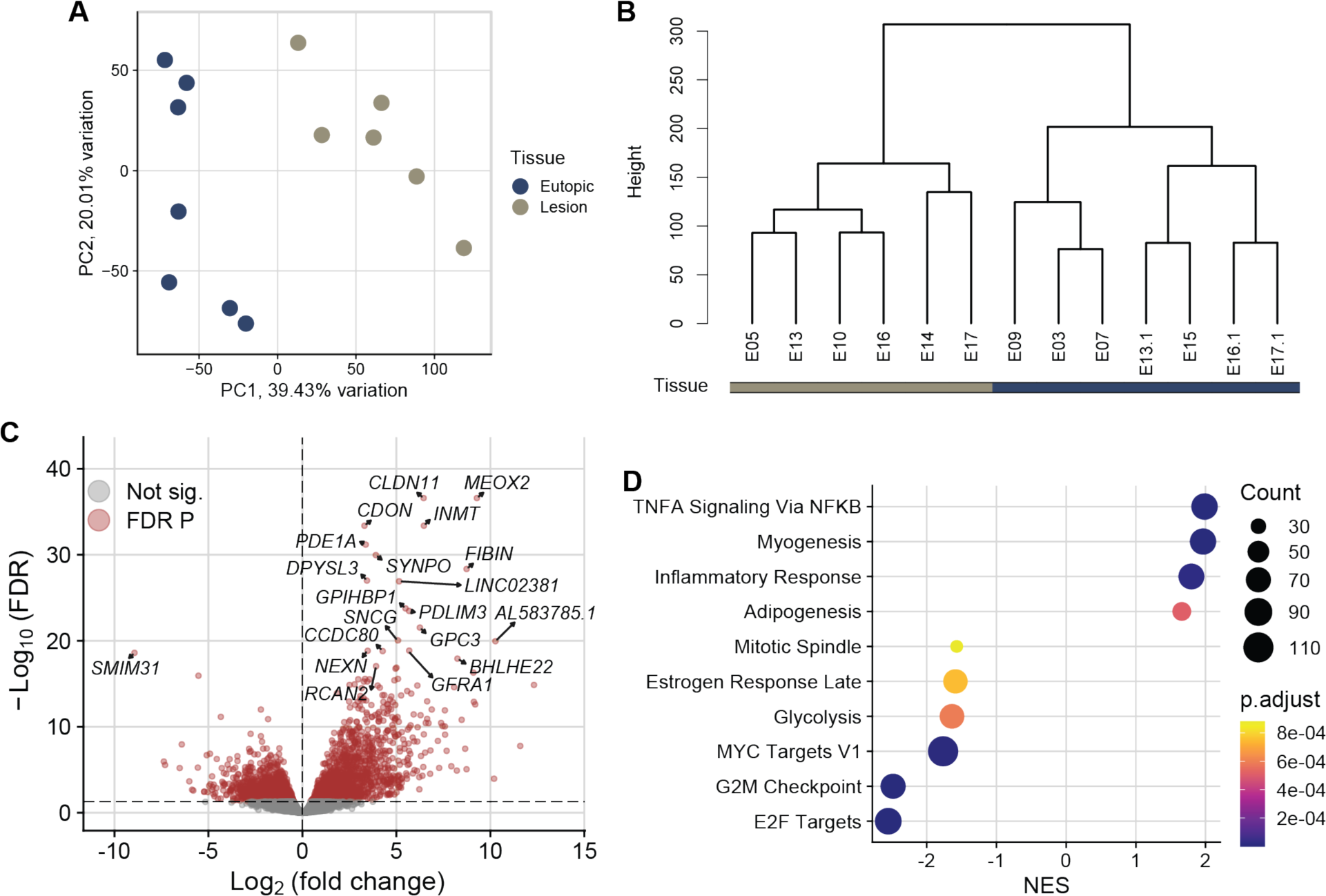
Transcriptome alterations in superficial endometriotic lesions compared to eutopic endometrium as measured by bulk RNA-sequencing. (A) Principal component (PC) plot of superficial peritoneal endometriotic lesions and eutopic endometrium (n = 13) from GSE179640. Samples were separated across PC1 by tissue type, representing 39% of variation in gene expression. (B) Hierarchical clustering dendrogram confirmed separation of lesions from eutopic endometrium. (C) Volcano plot of 3,656 differentially expressed genes in lesions versus endometrium. (D) Gene set enrichment analysis of hallmark pathways found inflammation, represented by TNF⍺ signaling via NF-κB and inflammatory response, increased in lesions while three cell cycle-related gene sets, MYC targets, G2M checkpoint, and E2F targets were decreased. This indicates increased inflammation and decreased proliferation in lesions compared to eutopic endometrium.

